# Heat induces multi-omic and phenotypic stress propagation in zebrafish embryos

**DOI:** 10.1101/2022.09.15.508176

**Authors:** Lauric Feugere, Adam Bates, Timothy Emagbetere, Emma Chapman, Linsey Malcolm, Kathleen Bulmer, Jörg Hardege, Pedro Beltran-Alvarez, Katharina C. Wollenberg Valero

## Abstract

Heat alters biology from molecular to ecological levels, but may also have unknown indirect effects. This includes the novel concept that animals exposed to abiotic stress can induce stress in naive receivers. Here, we provide a comprehensive picture of the molecular signatures of this process, by integrating multi-omic and phenotypic data. In individual zebrafish embryos, repeated heat peaks elicited both a molecular response and a burst of accelerated growth followed by a growth slow-down in concert with reduced responses to novel stimuli. Metabolomes of the media of heat treated vs. untreated embryos revealed candidate stress metabolites including sulphur-containing compounds and lipids. These stress metabolites elicited transcriptomic changes in naive receivers related to immune response, extracellular signalling, glycosaminoglycan/keratan sulphate, and lipid metabolism. Consequently, non heat-exposed receivers (exposed to stress metabolites only) experienced accelerated catch-up growth in concert with reduced swimming performance. The combination of heat and stress metabolites accelerated development the most, mediated by apelin signalling. Our results prove the concept of indirect heat-induced stress propagation towards naive receivers, inducing phenotypes comparable to those resulting from direct heat exposure, but utilising distinct molecular pathways. Group-exposing a non-laboratory zebrafish line, we independently confirm that the glycosaminoglycan biosynthesis-related gene *chs1*, and the mucus glycoprotein gene *prg4a*, functionally connected to the candidate stress metabolite classes sugars and phosphocholine, are differentially expressed in receivers. This hints at production of *Schreckstoff*-like cues in receivers, leading to further stress propagation within groups, which may have ecological and animal welfare implications for aquatic populations in a changing climate.

**Significance Statement:** Aquatic animals utilise chemicals to mediate adaptive behaviours. For instance, predated fish release chemical cues that elicit antipredatory responses in naive receivers. But whether abiotic factors such as heat likewise alter chemical communication has received little focus. Here, we uncover a novel dimension of chemical communication — heat-stressed donors can induce stress in naive receivers. We show that heat activates molecular stress responses, leading to the release of distinct stress metabolite classes into the environment. These stress metabolites alter the transcriptome of receivers, resulting in faster development and hypoactivity. Heat combined with stress metabolites had the largest effect, highlighting that abiotic stress, experienced both directly and indirectly, can alter chemical communication and affect embryonic development.

**Graphical Abstract:** 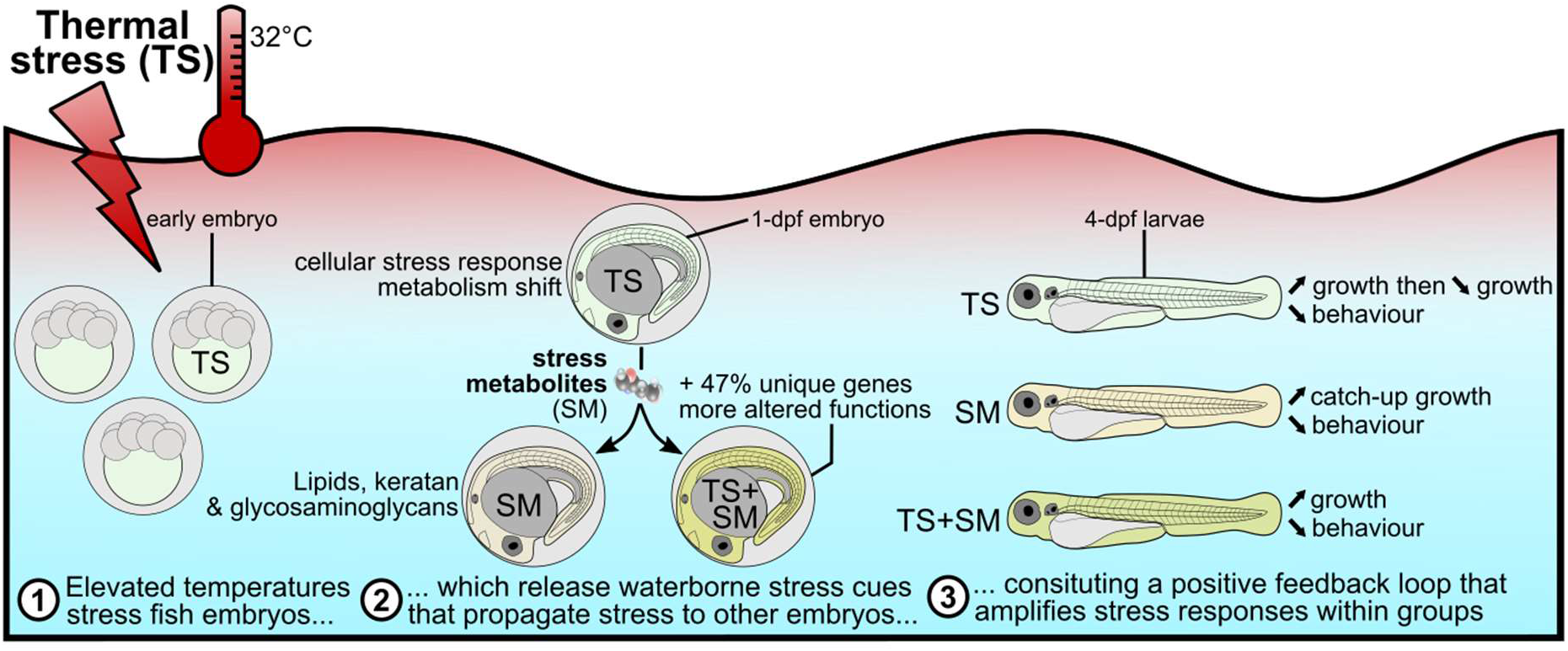

**Highlights:**

- We elucidate the mechanism for a novel dimension of the heat stress response — chemical communication from heat-stressed donors that induces stress in naive receivers — constituting a positive feedback loop
- Repeated heat stress induces a cellular and cortisol stress response and alters the phenotype of zebrafish embryos
- Heat-stressed embryos release stress metabolites enriched in lipids and sulphur-containing organo-oxygen compounds
- In combination, heat and stress metabolites induced 47% distinct differentially expressed genes, with many related to organ development
- These stress metabolites alter the transcriptome and induce both faster development and hypoactivity in naive receivers, a similar response to that of heat stress itself

## Introduction

Temperature is the abiotic “master” (1) factor regulating the biology of ectotherms (2). Heat alters a wide range of responses in fish, from gene expression (3) to development (4) and to population dynamics (5). The type of stimulus also matters: single and repeated periods of heat can induce different gene expression patterns leading to distinct behavioural stress responses (6). Repeated thermal conditioning can impair the ability to restore homeostasis (7) and alter the response to subsequent heat stress by attenuating the corticosteroid (8) and heat shock (9, 10) pathways. Furthermore, there is a growing concern about the response of fish embryos to heat due to their narrower thermal tolerance (11, 12), marking early development as a vulnerable “bottleneck” stage (13) when it comes to thermal stress. Heat stress responses at the molecular and behavioural levels are well studied in aquatic species, but its effect on chemical communication is less studied. Previous studies have focused mostly on cues released upon biotic factors such as injury (14) or disturbance (15). We have recently proposed that abiotic stress such as heat or low pH likewise elicits the release of olfactory cues, which we termed “stress metabolites” (16, 17). We found these cues to elicit similar phenotypic responses as heat stress itself in naive receivers, which are the characteristics of a positive feedback loop (16). To date, only a few studies have shown that animals communicate to each other upon abiotic stress, such as heat and low pH (17, 18).

Zebrafish (*Danio rerio*) is a commonly used experimental model species, including for behavioural assays (19) and “omics” approaches (20). Adult zebrafish tolerate water temperatures ranging 6-38°C (21) whilst zebrafish embryos develop normally in the 22-32°C range (22), with an optimum around 28.5°C (23). Importantly, thermal biology of zebrafish is conserved in laboratory conditions, which makes it a suitable model for investigating climate-related questions (24), such as the mechanistic basis of heat stress perception. Zebrafish embryos produce alarm cues (25) and innately react to extracts of crushed conspecific larvae with a decreased activity as early as 12-24 hours post fertilisation (hpf) (26). This early ability to detect cues is in line with observations of kin and alarm odour recognition in damselfish embryos (27). Altogether, this shows that fish, including zebrafish at early embryonic stages, can detect and discriminate stress chemical cues.

Natural chemical communication is likely mediated by a cocktail of compounds (28). However, stress-induced chemical communication has previously been studied using mainly phenotypic and physiological endpoints (15). Consequently, the biological compound bouquets mediating specifically heat stress-related communication, and their molecular pathways of action, remain unknown. Here, we explored the molecular response of zebrafish embryos exposed to thermal stress and to heat stress-induced metabolites using metabolomics and transcriptomics (Fig. 1, SI Appendix Fig. S1). We hypothesised that (i) the metabolomic profiles excreted by control and heat-stressed zebrafish embryos differ from each other; (ii) there may be similarities between the transcriptomic signatures of embryos exposed to heat stress and to stress metabolites, including in the response of chemosensory perception genes, and (iii) both heat-stressed and indirectly stressed individuals differ in growth patterns and behaviour from controls. (iv) Laboratory strains of zebrafish may perform differently to their wild counterparts (29), and embryos in clutches may react differently than single embryos in tubes. We expected to find similar molecular responses to stress metabolites in group-raised outcrossing zebrafish embryos, but perhaps with different magnitudes.

**Figure 1.**
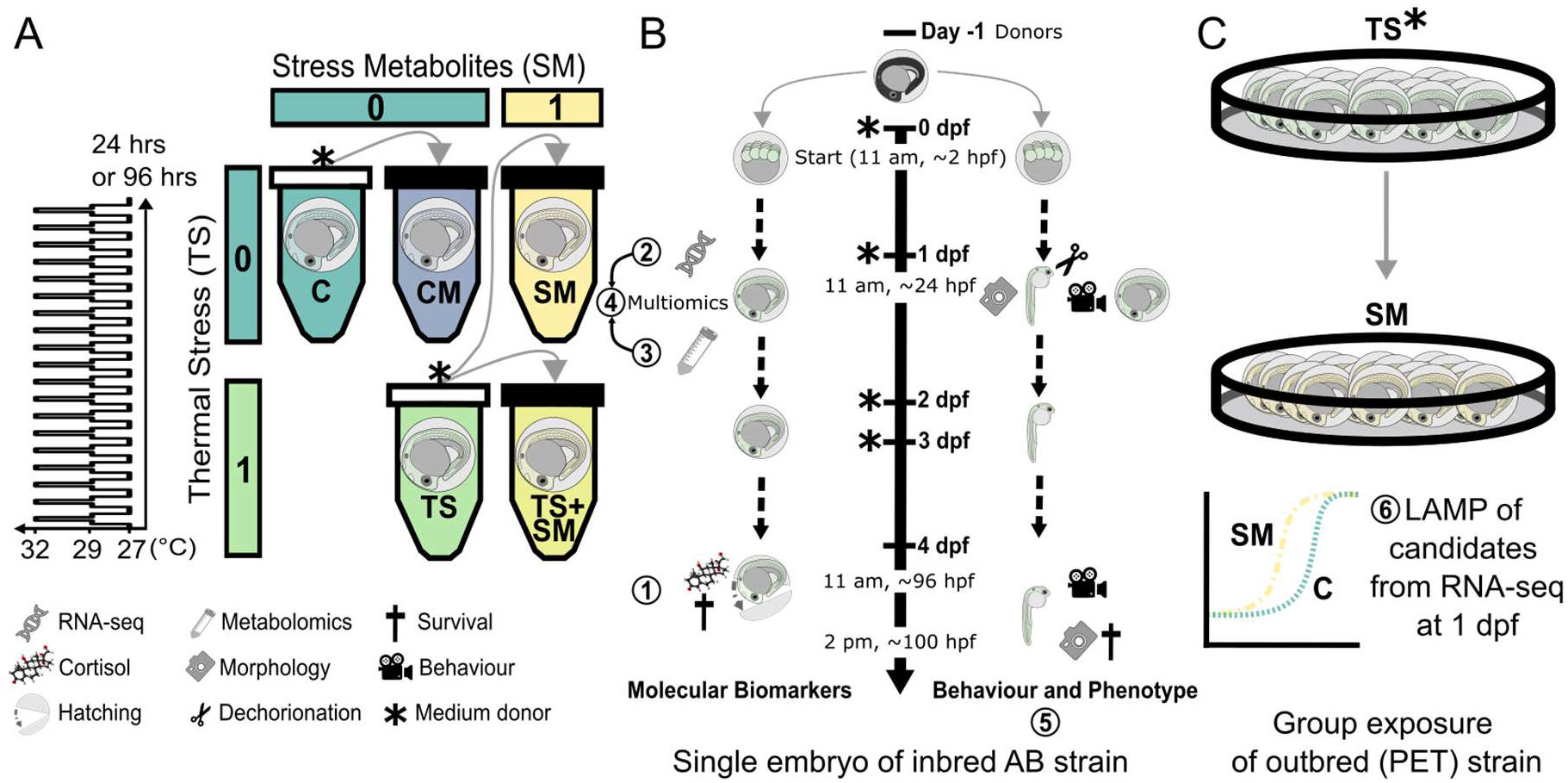
Scheme of experimental design. A) Zebrafish embryos (*Danio rerio*) < 3 hours post fertilisation (hpf) were incubated according to a two-way factorial design represented by two predictors: thermal stress (0 = control temperature of 27°C, 1 = repeated thermal stress at sublethal temperature, 32°C as shown in left graph) and stress metabolites (0 = fresh medium free of metabolites, 1 = stress metabolites released by embryos exposed to thermal stress). Treatments were CM: control metabolites at 27°C, C: control in fresh medium at 27°C, SM: stress metabolites at 27°C, TS: fresh medium in thermal stress, TS+SM: stress metabolites in thermal stress. Grey arrows indicate medium transfer from metabolite donor (✱ symbols) to metabolite receiver (plain black tube caps). B) Endpoints: embryos were sampled for both molecular (RNA-sequencing and metabolomics at 1 dpf and cortisol and HSP70 at 4 dpf) and phenotypic (1 dpf and 4 dpf) endpoints (circled numbers 1-5). C) Confirmatory experiment with Loop-Mediated Isothermal Amplification (LAMP) data of candidate genes found in RNA-seq in social groups of 20 embryos incubated in stress metabolites (SM) or fresh medium (C) until 1-dpf. Endpoints 1/2/5 utilised laboratory inbred AB and endpoints 3/6 wilder pet-store (PET) embryos.

## Materials and Methods

### Experimental design

Zebrafish embryos were collected by breeding adult zebrafish at the University of Hull. Embryos were collected in the morning, cleaned in fresh 1X E3 medium and rinsed by bleaching. Embryo stages were measured in hours post fertilisation (hpf) according to Kimmel et al. (1995) (23). Starting from the blastula period (~ 3.3 hpf), two temperature protocols were used, either (i) a control constant temperature of 27°C, or (ii) repeated temperature fluctuations from 27 to 32°C. A factorial design for the two factors “temperature” and “stress metabolites” (Fig. 1A), yielded five experimental treatments: fresh medium at 27°C (control C), fresh medium with thermal stress (TS), medium containing stress metabolites at 27°C (SM), and medium containing stress metabolites with thermal stress (TS+SM). A fifth condition containing medium collected from C contained control metabolites (CM). We measured five different endpoints at different time points, which yielded (i) cortisol, (ii) transcriptomic, (iii) metabolomics, (iv) multi-omic, and (v) phenotypic data (Fig. 1B). Additionally, we (vi) verified gene expression effects of SM in group-exposed, non-laboratory pet store (PET) line embryos (Fig. 1C). Refer to SI Appendix, SI Methods for further details.

### Endpoint 1: Cortisol and HSP70 response to heat and heat-induced metabolites in 4 dpf embryos

To determine the optimal time point to assess the cortisol stress response, we followed the dynamics of cortisol levels in control embryos at 1, 2, 3, and 4 dpf. For this purpose, control embryos were incubated in E3 medium in clutches at 27°C with natural (approx. 12:12 dark:light hours) light conditions. Embryos were pooled into groups of 40 embryos per sample (n = 3 pooled samples per time point). After optimisation for ELISA assay, zebrafish embryos were exposed in individual wells to C, CM, TS, and SM treatments from 0 to 4 dpf, until 60 embryos per biological replicate were obtained for each treatment (n = 3 samples per treatment, total of 180 embryos per treatment). Cortisol and HSP70 protein levels were then quantified in 4-dpf embryos.

### Endpoint 2: Transcriptomic response to heat and heat-induced metabolites in 1 dpf embryos

The transcriptomic response to thermal stress and stress metabolites was characterised by individually exposing zebrafish embryos to treatments C, TS, SM, and TS+SM for 24 hours. At the end of the exposure, viable zebrafish embryos still in their chorion were pooled into groups of 20 and processed for RNA-sequencing (n = 3 biological replicates of 20 embryos, i.e. total of 60 embryos per treatment). Reads were quality processed, annotated, and analysed for differentially expressed genes (DEGs), either as a complete dataset or by subset of candidate GO terms, and for functional enrichment of biological processes (BP), KEGG, and Reactome pathways. Since the zebrafish lateral line system is a candidate organ for chemical stimulus perception, we additionally compared the relative expression between hair cells and adjacent non-hair cells, of the SM DEGs vs. other genes expressed in zebrafish lateral line system hair cells from the dataset of Elkon et al. (66).

### Endpoint 3: Exploratory Metabolomics characterisation of heat-induced stress metabolites

We characterised metabolites released upon thermal stress. Zebrafish embryos were exposed for 24 hours. The protocol and medium samples resulted from our previous study (16) where the heating protocol consisted of thirteen heat peaks followed by a recovery period of 7 hours and 45 min at 27°C until 24 hours of treatment were reached, and medium would therefore not contain short-lived volatile cues. Media from donor embryos raised in C (i.e. control medium containing control metabolites CM) or TS (i.e. stress medium containing stress metabolites SM) were collected for metabolite identification. To account for contaminants in the E3 solution and microbial compounds, a blank was prepared by incubating fresh medium without embryos for 24 hours at 27°C. LC-MS/MS analysis was performed at the Metabolomics & Proteomics facility of the University of York. After data pre-processing and blank correction, masstags filtered for signal-to-noise threshold were retained as possible biomarkers of CM or SM (SI Appendix Fig. S1A). Biomarkers were compared using their unique m/z masses (± 0.0005 m/z) to known metabolites, including candidates (SI Appendix Table S1), from publicly available metabolomics databases. Biomarkers of either CM and SM groups were then functionally enriched for KEGG pathways and chemical subclasses.

### Endpoint 4: Multi-omic analysis

To better connect the genetic responses to heat stress by donors secreted metabolites with the response to these stress metabolites by receivers, transcriptomic and metabolomic data were integrated into joint analyses. using Joint Pathway analyses and compound/protein interactions (CPI). We reasoned that DEGs in TS may lead to the synthesis and release of stress metabolites which would initiate a transcriptome response in receiver embryos. Hence, both the joint pathway and CPI analyses were performed by combining the stress metabolites with DEGs of either SM or TS.

### Endpoint 5: Phenotypic response to heat and heat-induced metabolites in 1 dpf and 4 dpf embryos

A range of phenotypic parameters was measured in zebrafish embryos at 1 dpf and 4 dpf in all five treatments, both within and outside of their chorions after dechorionation. All conditions were tested twice on two independent batches at different dates, using each time one embryo clutch for all treatments to limit batch effects. At 1 dpf, embryos were imaged (SI Appendix Fig. S1B) and light-induced startle responses were recorded. At 4 dpf, larvae were imaged, and touch-evoked behaviours were videoed. Images and videos were randomised for blind analysis.

### Endpoint 6: LAMP characterisation of candidate genes in response to heat stress metabolites in outcrossing group-exposed embryos

Outcrossing pet store line (PET) zebrafish were exposed in groups (3 sets of 20 embryos each) in petri dishes for 24 hrs to either fresh E3 medium or stress metabolites (n = 60 embryos per sample, 250 μL medium per embryo). Stress metabolites were obtained from donors experiencing constant thermal stress of 32°C from 0 to 1 dpf. Total RNA was extracted from 4 such pools per treatment as previously described to quantify *chs1*, *ldha*, *ora3*, *otofa*, *prg4a*, and *tlr18* using fluorimetric LAMP technology.

## Results

### Heat stress, but not stress metabolites, leads to increased cortisol in 4-dpf larvae

Cortisol plays a central role in the fish hypothalamic-pituitary-interrenal (HPI)-axis stress response. We first confirmed that cortisol increased between 1 and 4 dpf in control embryos (SI Appendix Fig. S2A-B). Cortisol was then measured at 4 dpf in unheated control (C) and heated (TS) medium donors and their respective receivers of media CM (control metabolites) and SM (stress metabolites) (SI Appendix Fig. S2C, Table S2). A one-way ANOVA showed that cortisol concentration significantly varied across treatments (F = 14.35, P = 0.0014). Post-hoc pairwise comparisons revealed strong evidence that TS significantly increased cortisol concentrations by 92% compared to control C (t = −4.86, P = 0.0055, huge effect size = 6.17), but incubation media did not have an effect (SI Appendix Table S2). In contrast to cortisol, protein levels of HSP70 were similar across all treatments at 4 dpf (F = 1.9271, p = 0.2038, SI Appendix Fig. S2D).

### Heat stress alters the whole-body transcriptome of 1-dpf embryos

Repeated heat stress induced n = 369 DEGs (p ≤ 0.01) (Fig. 2A, and PCA, SI Appendix Fig. S3). Functional enrichments (BP, KEGG, Reactome) were conducted for the top 106 genes (Dataset S1) with strongest support (p-adj ≤ 0.05), most of which were upregulated (n = 90 genes with fold change FC > 1.5) whilst a few were downregulated (n = 16 genes). The three genes with largest effect size and log fold-change |LFC| > 3 were *atp2a1l*, *crygm2d18*, and *matn3a*. Two paralogues of gamma-glutamylamine cyclotransferase, tandem duplicate (*ggac*t.2 and *ggact.3*), involved in glutathione metabolism, were the two most heat-inhibited genes, with 98% fewer transcripts than in control embryos. Heat stress induced transcriptomic changes in embryonic development (eyes, muscles, and somites), in the heat stress response, and the metabolism of sugars, amino acids, purine, and energy intermediates (ATP and pyruvate, Fig. 2D, SI Appendix Supplementary Results, Fig. S4, Dataset S1). In addition, network analysis revealed strong functional interconnectivity between the DEGs of TS (Fig. 3A, CPI enrichment p < 0.0001, 64 edges observed vs. 5 expected edges). A cluster of co-upregulated genes was assigned to the GO term “metabolism of carbohydrates” (GO:0005975): *pgk1, gpia, eno1a, eno3, ldha*, and *gapdhs*. Similarly, another cluster grouped five transcription regulators (GO:0000122) of the GO terms “segmentation” (GO:0035282) and “somite development” (GO:0061053): *mespab, her1/7, ripply2*, and *tbx6* (Fig. 3A, Dataset S1).

**Figure 2.**
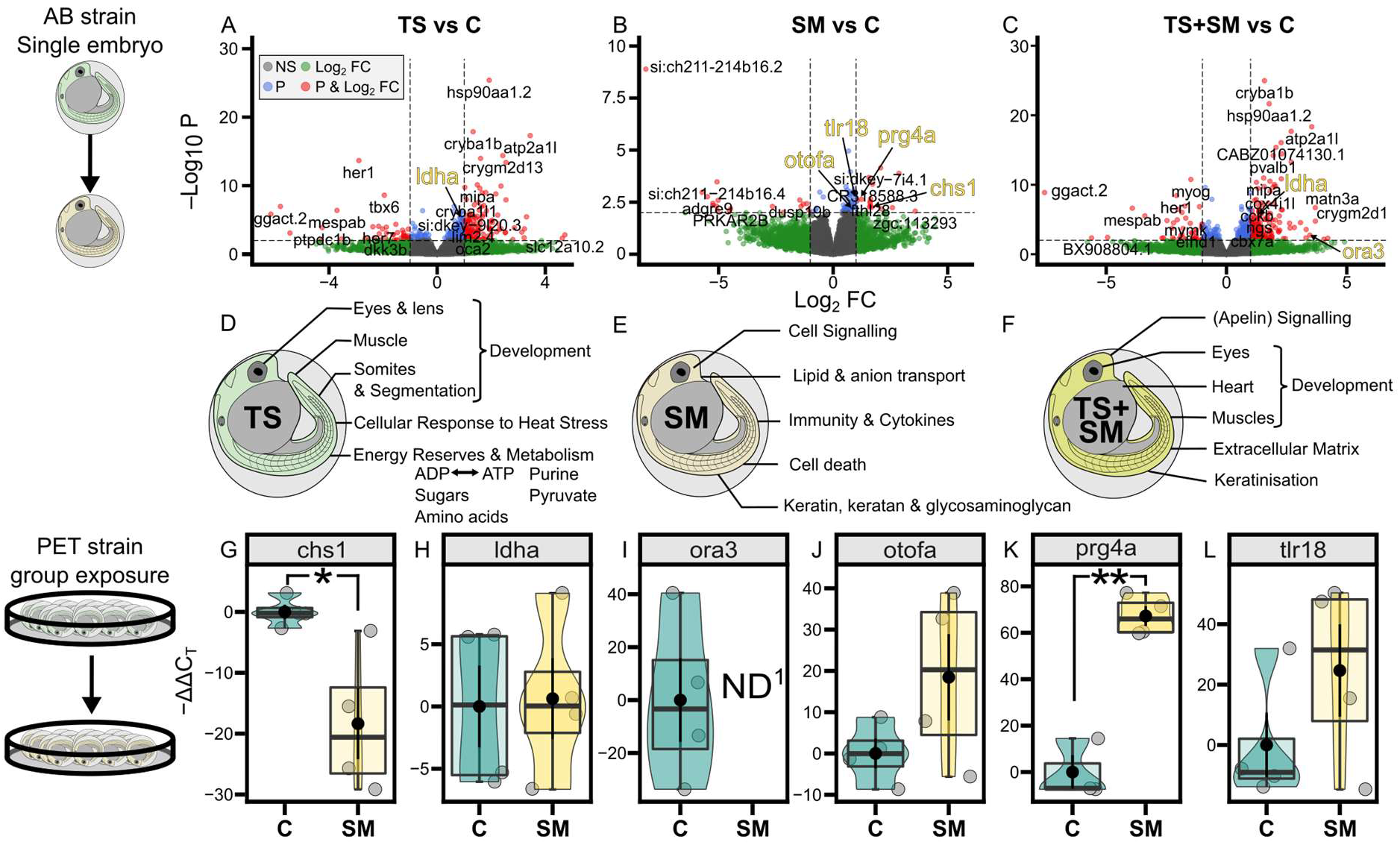
Thermal stress and Stress Metabolites alter the transcriptome and its functions in 1-dpf zebrafish embryos. Top row: volcano plots showing the differentially expressed genes in response to (A) thermal stress (TS), (B) stress metabolites (SM), and (C) their combination TS+SM compared to control C. Genes of interest are shown in red when they have significant raw p-values (above horizontal line) and an absolute fold change (FC, representing the effect size) greater than 2 (|log 2 FC| > 1, vertical lines). DEGs left to the left vertical line and right to the right vertical line are respectively significantly underexpressed and overexpressed compared to the control C. The dotted arrow in B) shows the significant gene in SM after p-value adjustment. Middle row: gene functional categories from KEGG, Reactome, and GO Biological Process analysis for (D) TS, (E) SM, and (F) those uniquely present in the combined treatment TS+SM. Top and middle row show transcriptomic data from individually-raised AB strain embryos. Bottom row shows Loop-Mediated Isothermal Amplification (LAMP) data of candidate genes associated with stress metabolites showing the normalised time of detection (-ΔΔCT) in C and SM in more realistic environmental conditions (genetically diversified outcrossing pet-store PET strain raised in groups). *chs1*: chitin synthase 1, *ldha*: lactate dehydrogenase A4, *ora3*: olfactory receptor class A related 3, *otofa*: otoferlin a, *prg4a*: proteoglycan 4a, *tlr18*: toll-like receptor 18. ^1^ND: mRNA amplification non-detected suggesting a depletion of *ora3* in SM. Student’s t-tests compared SM to C with significant comparisons shown by horizontal bars with *: p ≤ 0.05, **: p ≤ 0.01. Treatments were SM: stress metabolites at 27°C, TS: fresh medium in thermal stress, and TS+SM: stress metabolites in thermal stress, compared to C: control in fresh medium at 27°C.

**Figure 3.**
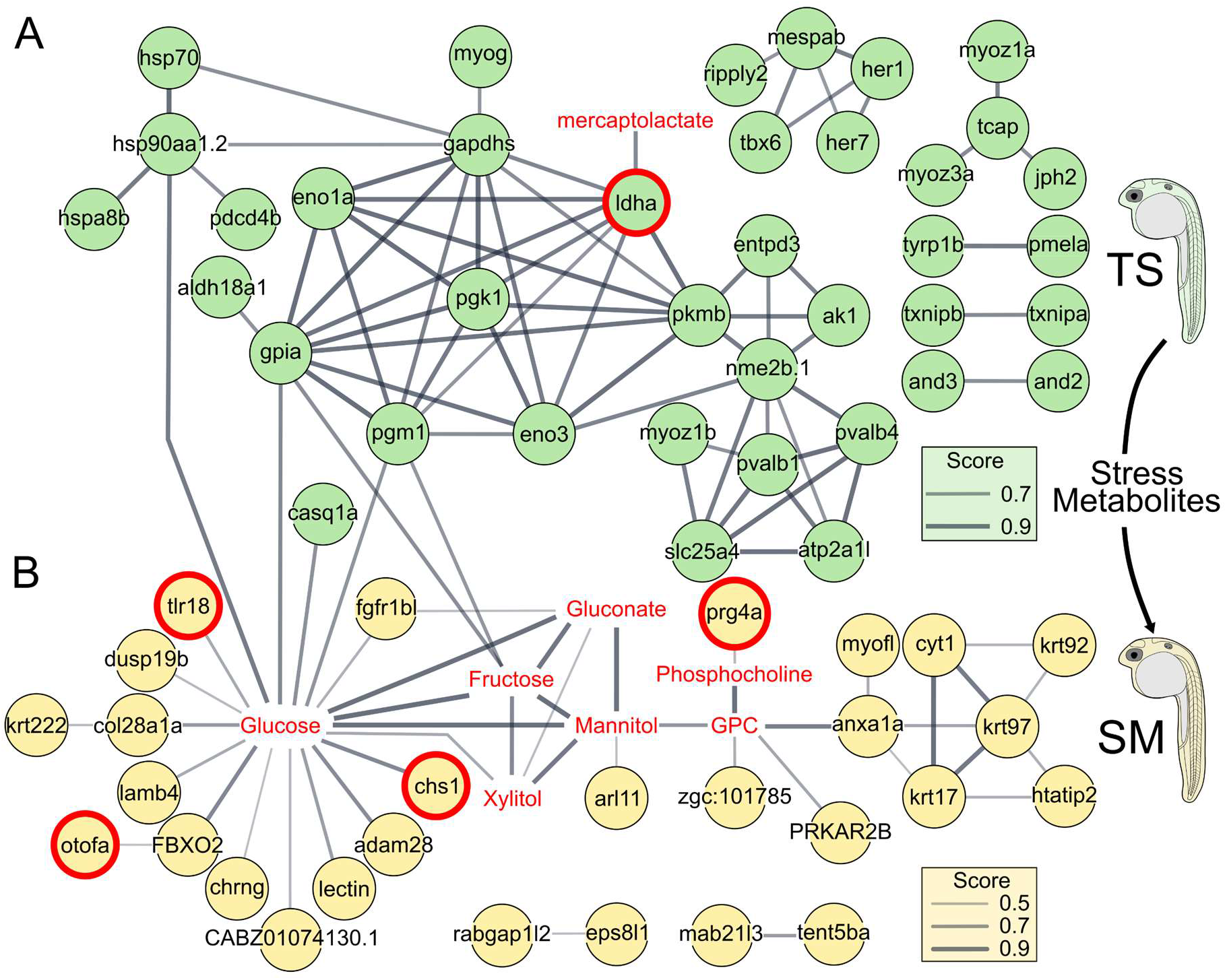
Heat-induced DEGs are functionally connected to stress metabolites, which interact with the transcriptome of receivers. Compound-Protein Interaction (CPI) network analysis of significant differentially expressed genes (circles) of TS (top row, A) or genes of SM (bottom row, B) and stress metabolite compounds (in red without circle). Line width represents the increasing confidence score with a minimum threshold of 0.7 (TS) or 0.4 (SM). Computed and drawn independently for A and B in STITCH using Cytoscape and merged in Inkscape. A: CPI enrichment P < 0.0001, 64 edges observed vs. 5 expected edges. B: CPI enrichment p < 0.0001, 14 CPI edges observed vs. 3 expected. GPC: glycerophosphocholine. Treatments were SM: stress metabolites at 27°C and TS: fresh medium in thermal stress compared to C: control in fresh medium at 27°C. Genes with red bold circles were used for LAMP measurements.

### Stress metabolites induce a localised transcriptomic signature in 1-dpf embryos

While the transcriptomic response to the SM treatment was less pronounced compared to the whole-body response to heat stress, stress metabolites altered 79 transcripts with raw p-values lower than 0.01 (Fig. 3B, Dataset S1). Respectively, 45 and 15 of these were up- or downregulated in SM compared to C. The gene *si:ch211-214b16.2* (ENSDARG00000102593, significant after p-value adjustment) was downregulated by 99.7% by stress metabolites and is likely an ortholog of the gene NOD2 identified as orthogroup *45875at7898* at Actinopterygii level by OrthoDB v10.1 (30). Other than expected, genes belonging to the GO term “chemosensory perception” (GO:0007606) were not significantly enriched (Dataset S1). Instead, the transcriptomic response to SM was related to four wider categories: extracellular signalling, (phospho-) lipid and anion transport, the metabolism of glycosaminoglycan and keratan, but also the immune response involving interleukin cytokines and apoptosis (Fig. 3B, SI Appendix Supplementary Results, Fig. S5A-C, Dataset S1). The glycosaminoglycan/keratan/keratin term involved several genes including *krt17, krt97*, chitin synthase 1 (*chs1*), carbohydrate (N-acetylglucosamine-6-O) sulfotransferase 2b (*chst2b*) but also a proteoglycan *prg4a*. SM-responsive genes were also significantly (p < 0.0001, 14 CPI edges observed vs. 3 expected) functionally connected in a network (Fig. 3B) which supports the biological relevance of the measured response. Remarkably, the DEGs in SM are also significantly upregulated in zebrafish lateral line hair cells in 5 dpf larvae, relative to their expression in neighbouring epidermis cells (z-score = 8.71505; P < 0.0001, SI Appendix Fig. S5D). One interesting gene emerging from this comparison was *xkr8.2* (ENSDARG00000076820), which is the highest upregulated gene in both hair cells and SM (LFC = 1.68) compared to non-hair cells, and is a scramblase involved in the enriched GO term “membrane phospholipid transport” (GO:0015914, Dataset S1).

### Heat stress and stress metabolites induce few similar genes and have unique effects in combination

Nine genes were common to both individual TS and SM conditions compared to control C. Genes upregulated in both conditions were associated with eye development (*cryba1b*), the immune response (*tcima*), chloride transmembrane transport (*best1*), and cell death (*fthl28*). In contrast, *ggact.2*, involved in glutathione metabolism, was inhibited by TS but upregulated in SM. In the natural environment, if heat stress occurs, SM would increase simultaneously. We therefore explored the transcriptome of the combined TS+SM treatment, which was mainly driven by heat stress as 49% DEGs (78/159) were shared between TS and TS+SM (Fig. 2C, SI Appendix Fig. S6A), with functions being largely similar to those in TS (SI Appendix Supplementary Results, Figs. S6B, S7A, Dataset S1). However, the combined treatment also up-regulated one olfactory gene, the taste receptor *ora3* (olfactory receptor class A related 3, GO term-wide p-adj = 0.0412, Dataset S1). Moreover, 47% of DEGs (75/159 with p-adj < 0.05) were uniquely up- or down-regulated in TS+SM condition, suggesting an interaction between the SM and TS factors causing a different response than either factor by itself, with additional functions unique to TS+SM ascribed to developmental processes (eyes, heart, muscles). Interestingly, this involved apelin signalling (with five genes *acta2*, *si:ch211-286b5.5, mef2aa, ryr2a, map1lc3cl*) (Fig. 2F, SI Appendix Fig. S6C-E, Dataset S1). We additionally compared the treatment TS+SM to SM and TS to determine responses of combined *versus* single stressors (SI Appendix Fig. S7B-C, Dataset S1). One particular gene that was significantly altered by TS+SM compared to SM was *olfcb1*, (olfactory receptor C family, b1, LFC = 2.30, p-adj < 0.0001) suggesting that heat in combination with SM affects chemosensation. Comparing TS+SM to the TS treatment showed that the addition of SM adds 32.5% novel BP terms, 37.5% Reactome pathways, and 16.7% KEGG pathways (SI Appendix Fig. S7D-E).

### Stress and control media show distinct metabolomic profiles

The metabolomic profile of the stress metabolites contained 89 metabolites which were matched to 658 possible compounds (Fig. 4A, SI Appendix Fig. S8, Dataset S2). Correlating masstag intensities in SM and CM showed that metabolites were either present in both media in varying concentrations, or were uniquely secreted in either CM or SM (dubbed “CM cloud” and “SM cloud” Fig. 4A, SI Appendix Table S3). Two different masstags were assigned to 3’-mercaptolactate (SM12 and SM21). Several a priori defined candidate chemicals for stress metabolites (i.e. from alarm/disturbance cue studies, SI Appendix Table S1) were unexpectedly more concentrated in control medium compared to stress medium, such as hypoxanthine-3-N-oxide (C5H4N4O2, for which we found hypoxanthine C5H4N4O), trigonelline or homarine, cysteine, or acetylcholine (SI Appendix Table S3, Dataset S2). Representative compounds of each identified masstag were used for functional subclass and KEGG pathway enrichments (SI Appendix Fig. S9, Dataset S2), which revealed that control metabolites were mainly associated with amino acids (e.g. CM8 proline, CM14 L-tyrosine), catechols, purine ribonucleoside monophosphate (e.g. CM6 adenosine monophosphate), and dipeptides. Stress metabolites in contrast were mostly associated with sulphur-containing carbothioic S-acids (organosulfur compounds), and lipids including O-acylglycerol-phosphates, fatty acids, and glycerophospholipids (SI Appendix Fig. S9). The representative stress metabolite compounds were mainly ascribed to lipid molecules and “organic oxygen compounds’’, whereas control medium contained molecules which had a more diverse chemical classification and were mainly “organic acids and derivatives’’ and “nucleosides, nucleotides, and analogues” (Fig. 4B-C, SI Appendix Table S3, Dataset S2). Several compounds did not have ChemOnt classification but were likely proteins (SM1, SM4, SM19, SM23) or polyamines (SM7).

**Figure 4.**
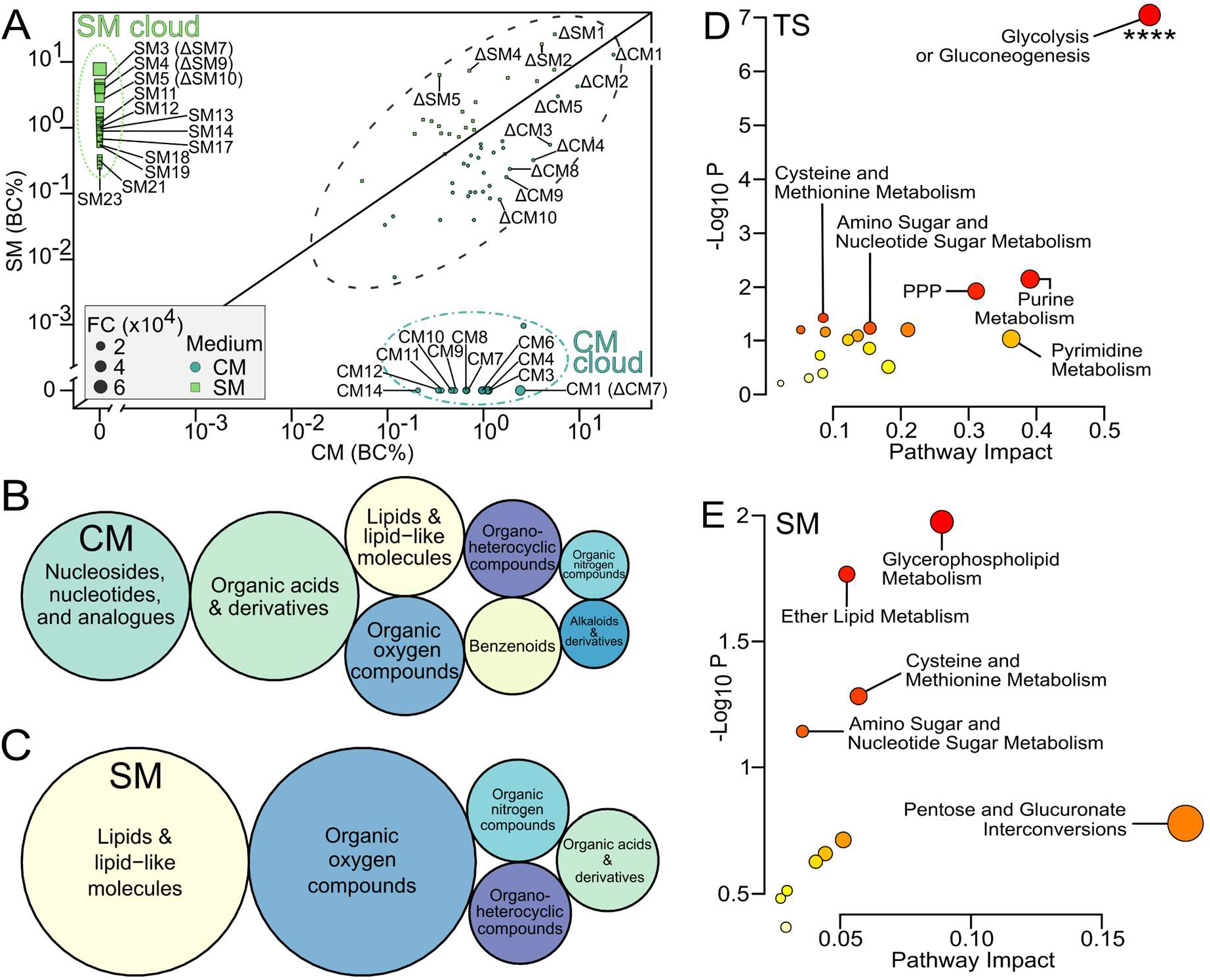
Multi-omic data evidence that stress metabolites differ in classes and functions from control metabolites. A) Correlation plot showing filtered (n = 89) masstags that are possible biomarkers of control metabolites CM (blue circles, bottom right) and stress metabolites SM (green squares, top left) groups. Data is presented as blank corrected total percent intensity (BC%). Masstag inside the black dotted ellipse are unlikely to be biomarkers of SM or CM since they are present in both conditions. Masstags inside the dot-dashed blue and dashed green ellipses represent CM and SM-specific biomarkers, respectively. FC: fold-change between SM and CM where the numerator is the medium with the highest concentration for each compound. Labels show the masstags with possible hits amongst the top 23 compounds of SM and top 14 compounds of CM based on fold-change (see Table S3 for associated chemicals) and top 10 compounds based on difference between SM and CM (delta Δ identifiers). Superclasses of representative hits for the biomarkers of (B) CM and (C) SM. Bubble sizes are proportional to counts of superclasses per medium type. Candidate cause-and-effect pathways derived from multi-omic analysis of KEGG pathways integrating the representative hits of the stress metabolites with the differentially expressed genes of (D) TS or (E) SM. The top five most evident pathways are given within plots, respectively. Significant terms are shown by **** (p-adj ≤ 0.0001). PPP: pentose phosphate pathway. See Dataset S2 for class-sorted compounds.

### Multi-omic analysis uncovers sender-receiver gene-metabolite candidate cause and effect pathways

Since (i) the DEGs of heat-stressed donors (TS vs C) putatively lead to the synthesis and/or release of stress metabolites (SM), which (ii) may in turn have regulated the transcriptomic signature in receivers (DEGs from SM vs C), we explored this possibility using joint pathway and STITCH analyses (Dataset S2). The joint pathway analysis evidenced that donors significantly enriched the “glycolysis/gluconeogenesis” pathway and compounds. (Fig. 4D). Receivers, in contrast, enriched, albeit with weak evidence (raw p ≤ 0.07 but FDR > 0.05), glycerophospholipid, ether lipid, cysteine and methionine, and amino sugar metabolism pathways (Fig. 4E, Dataset S2). The compound-protein interaction network evidenced a strong connectivity of stress metabolites with DEGs of either TS (64 CPI edges observed vs. 5 expected edges, p < 0.0001) or SM (14 CPI edges observed vs. 3 expected) with stress metabolites (Fig. 3). For instance, *ldha* (Lactate dehydrogenase A4) was linked to 3’-mercaptolactate in donors (Fig. 3A) whereas in receivers many genes, including *tlr18* (toll-like receptor 18), *chs1* (chitin synthase 1), and *otofa* (otoferlin A), linked to glucose, whilst *prg4a* (proteoglycan 4a) was associated with phosphocholine (Fig. 3B).

### Both heat stress and heat-induced stress metabolites alter development and stimulus-induced behaviour

Heat stress (F = 30.1, R^2^ = 0.25, P = 0.001), but neither stress metabolites (F = 0.16, P = 0.7610) nor the interaction term (F = 2.34, P = 0.1320), altered the phenotype of 1-dpf embryos (Fig. 5A, SI Appendix Table S4). Post-hoc tests confirmed that thermal stress significantly altered embryo growth, reaching the pharyngula stage earlier (z = −2.67, P = 0.0076, Fig. 5C, SI Appendix Fig. S10B). Conversely, stress metabolites had no effect on growth at 1-dpf. However, 4-dpf zebrafish embryos grew significantly longer in the presence of stress metabolites (t = 4.80, P = 0.0304, small effect size = 0.35) but not under thermal stress (t = 1.91, P = 0.1692) nor the interaction term (t = 0.11, P = 0.7353, Fig. 5D). An ANOVA revealed that both heat and stress metabolites significantly altered the increment in growth (ΔSEL) between 1 and 4 dpf, but in opposite directions (Fig. 5E, SI Appendix Tables S5-S6). Heat-exposed embryos had a lower ΔSEL (t = 20.99, P < 0.0001, large effect size = 0.81) whereas those experiencing stress metabolites accelerated growth from 1 to 4 dpf (t = 6.65, P = 0.0112, small effect size = 0.43) and slightly surpassed the TS treatment in average final length at 4 dpf (Fig. 5D-E). Heat stress (t = 18.15, P < 0.0001), but neither stress metabolites (t = 0.29, P = 0.5891) nor their interaction (t = 0.07, P = 0.7872), significantly reduced the startle response of 1-dpf zebrafish embryos (Fig. 5B, SI Appendix Tables S7-8). Heat also slowed mean acceleration (t = 4.26, P = 0.0412, small effect size = 0.34) and TS marginally reduced mean speed compared to C (t = −2.74, P = 0.0212) in the touch-evoked swimming response of 4-dpf larvae (SI Appendix Fig. S11B-C). Larvae incubated in stress metabolites, however, swam fewer burst counts compared to fresh medium (t = −2.1, P = 0.0381, Fig. 5F). See SI Appendix Supplementary Results for an extensive description of phenotypic data.

**Figure 5.**
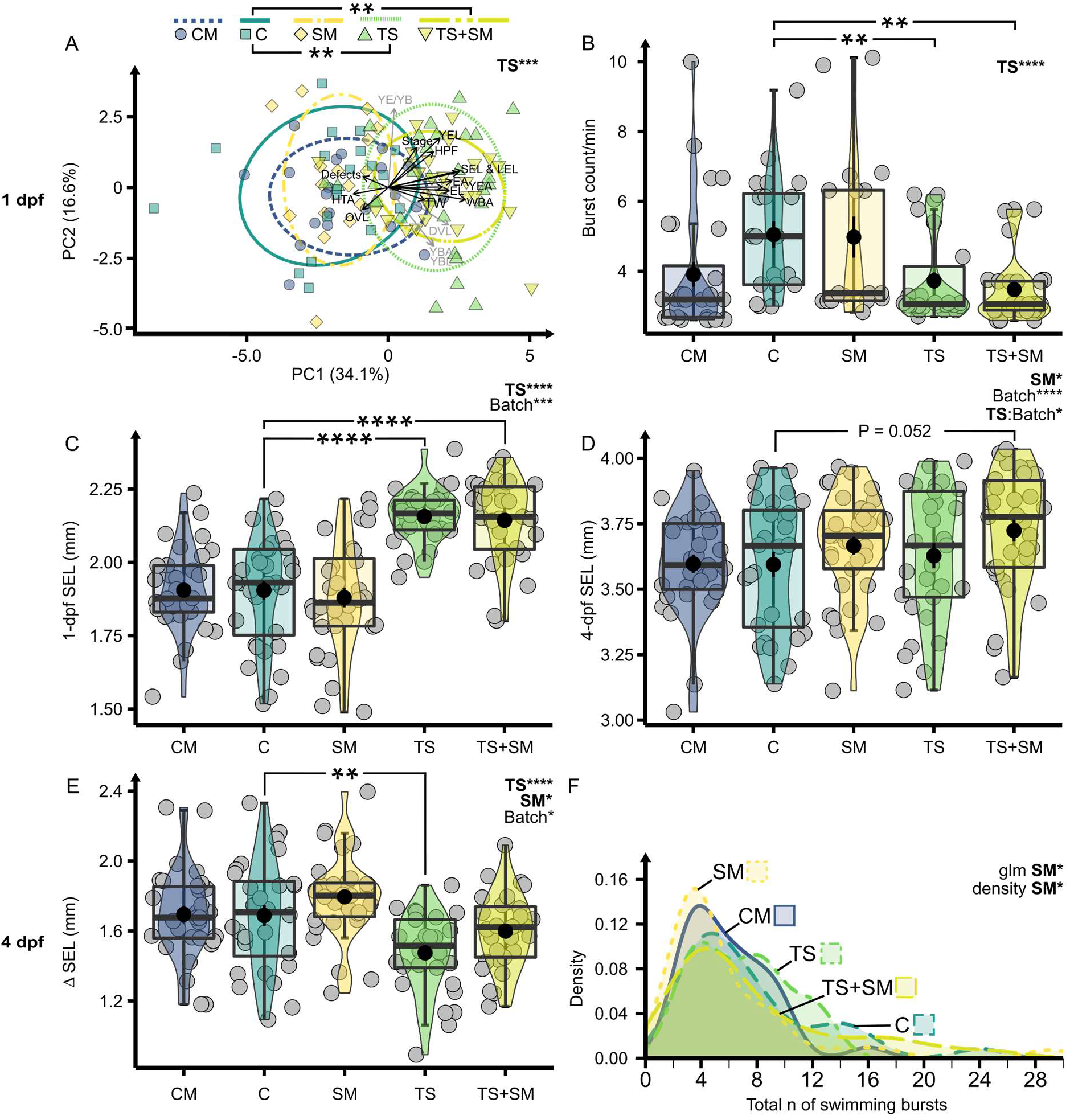
Thermal stress and stress metabolites alter the development and behaviour of zebrafish embryos. A) PCA of 1-dpf morphology. HPF: final stage in hours post fertilisation, Stage: % in segmentation or pharyngula, EA: eye area, EL: eye length, DVL: dorsal-ventral length, HTA: head-trunk angle, LEL: longest embryo length, OVL: otic vesicle length, SEL: shortest embryo length, WBA: whole-body area, YBA/L: yolk ball area/length, YEA/L: yolk extension area/length, YE/YB: yolk extension/yolk ball ratio. Individual variables significantly altered by thermal stress are shown by black arrows. B) Burst count per minute in active 1-dpf embryos. C) SEL (mm) increment from 1 to 4 dpf. D) Mean acceleration (m/s^2^) of the 1^st^ complete burst following 1^st^ and 3^rd^ touch stimuli. E) Density distribution of total burst count and total distance in cm (F) after three touch stimuli. Effects of thermal stress and stress metabolites across the two-way factorial design (C, SM, TS, TS+SM) were computed using PERMANOVA (A), ANOVA (B-D, F), or negative binomial generalised linear model (E, confirmed by a density test) with significant predictors and covariates (“batch”) reported on top-right corners. Pairwise tests with FDR p-value correction compared CM to C and SM, or C to SM, TS, and TS+SM, with significant comparisons shown by horizontal bars. *: p ≤ 0.05, **: p ≤ 0.01, ***: p ≤ 0.001, ****: p ≤ 0.0001. Treatments were CM: control metabolites at 27°C, C: control in fresh medium at 27°C, SM: stress metabolites at 27°C, TS: fresh medium in thermal stress, TS+SM: stress metabolites in thermal stress.

### Stress metabolites also exist in group-raised non-laboratory embryos

In our laboratory AB line experiments, we found a localised response to stress metabolites and a lower phenotypic response than we found previously using PET line embryos (SI Appendix Fig. S12, (16)). An independent confirmatory experiment utilising non-laboratory (PET line), group-raised zebrafish embryos therefore aimed to confirm the role of candidate genes from the RNA-seq data in the response to stress metabolites. First, *ldha*, overexpressed in both TS and TS+SM in the RNA-seq data, was not altered in response to stress metabolites (t = −0.14, p = 0.8960, Fig. 2H, Table S9). Two candidate genes involved in sensory perceptions (*otofa*, expressed in lateral line hair cells) and immunity (*tlr18*) were (non-significantly) upregulated by stress metabolites (Fig. 2J, L, Table S9). Stress metabolites significantly upregulated *prg4a* (t = −8.01, p = 0.0026, |d| = 6.52, “huge” effect size), with all of these genes’ expression patterns matching the RNA-seq data of the AB line zebrafish. In contrast, chs1 was significantly downregulated (t = 3.08, p = 0.0487, |d| = 2.18, “huge” effect size, Fig. 2G, K, SI Appendix Table S9) in this experiment, contrasting with significant upregulation in SM of the AB line experiment (LFC = 1.64). One chemosensory candidate gene, *ora3* (olfactory receptor class A related 3), could not be detected in the stress metabolite treatment, suggesting receptor depletion in embryos exposed to stress metabolites (Fig. 2I).

## Discussion

The aim of this work was to elucidate the molecular aspects of heat-induced stress propagation in zebrafish embryos. We first confirmed that repeated heat peaks alter a range of different aspects of the biology of zebrafish embryos. In our study, heat stress altered molecular profiles leading to developmental changes, and earlier hatching. Contrasting with studies reporting hyperactivity upon heat stress (31, 32), zebrafish embryos were hypoactive, which was mirrored by transcriptomic enrichments of muscle genes and functions. We confirmed the cortisol stress response onset around 2-4 dpf in zebrafish (33, 34). The central role of glucocorticoids in muscle growth, hatching, and behaviour (35, 36) may explain the concomitant increase in cortisol and the decrease in locomotor activity observed here. The upregulation of heat shock protein (HSPs) transcripts and cortisol confirmed that repeated heat stress induced a (cellular and heat) stress response (37, 38). Similarly, overregulated *hsp90a* transcripts were found in marine medaka *O. latipes* upon repeated thermal stress (39). Heat stress induced glucose, pyruvate, and ADP/ATP metabolism, indicating mobilisation of energy reserves. Heat stress altered developmental transcriptome dynamics. As a consequence, heat-stressed embryos were older and grew longer at 1-dpf, in accordance with previous studies (4, 16, 40). However, we observed a subsequent slow-down of development from 1 to 4 dpf conforming to the temperature-size rule stating that fish growing faster during early stages become smaller adults (41).

Only a few prior studies have characterised stress-induced whole-body metabolome profiles in fish (42–44), but none yet have focused on cue secretion into the environment. A major finding in our study was that heat induces the release of stress metabolites into the medium that are distinct from control metabolites released by unstressed embryos. The multi-omic compound-protein interactome evidenced functional links between transcriptomes and the metabolome containing stress metabolites released into the medium. Interestingly, several candidate chemicals (e.g. trigonelline, homarine, hypoxanthine-3-N-oxide) previously identified as alarm cues (45, 46) were more concentrated in the control medium. In contrast, stress metabolites were mainly ascribed to two compound superclasses: “organic oxygen compounds” and “lipids and lipid-like molecules”.

Carbohydrates within stress metabolites may originate from heat-induced metabolic activity such as glycolysis (47). The multi-omic analysis highlighted that the pentose phosphate pathway (PPP, including deoxyribose, *gpia*, and *pgm1*), a major cellular redox mechanism (48), was altered in response to heat. Organosulfur compounds (such as 3-mercaptolactate or 2-oxo-4-methylthiobutanoic acid) were a main component of the organic oxygen compounds classification. Multi-omic analysis of stress metabolites with receiver DEGs revealed “Cysteine and Methionine metabolism” as the third most important altered pathway, lending support to a functional relationship between secreted organosulfur compounds and receiver responses. 3-mercaptolactate is part of the cysteine transamination pathway and synthesised by the heat stress-induced DEG lactate dehydrogenase A *ldha* (49). Lactate dehydrogenase was likewise found to be upregulated by heat stress in rockfish *Sebastes schlegelii* (42). Organosulfur compounds may function as cues in chemical ecology, for instance in fox urine 50 and cat urinary pheromones (51), and therefore warrant further study.

In addition, stress metabolites predominantly contained a range of lipids, including fatty acyls, glycerolipids, sphingolipids, glycerophospholipids, and steroid esters. The multi-omic analysis found “glycerophospholipid metabolism” and “ether lipid metabolism” as the two major KEGG pathways enriched between stress metabolites and receiver transcriptomes. Previous lipidomic studies demonstrated a role of lipids, particularly sphingolipids, in protecting the cell from heat damage and acting as messengers (52, 53). Differential vulnerability of lipid classes to reactive oxygen species leads to temperature-controlled membrane lipid remodelling (54, 55). For instance, the proteome of adult zebrafish shows such changes in lipids following heat stress at 34°C for 21 days (56) and crucian carps (*Carassius carassius*) release phosphatidylenolamines upon alkalinity stress (44). Two notable glycerophospholipids amongst stress metabolites were phosphocholine and glycerophosphocholine. Olive Flounder *Paralichthys olivaceus* exposed to heat stress also displayed elevated tissue levels of phosphocholine and glycerophosphocholine (57), which may originate from temperature-induced breakdown of phosphatidylcholine in the cell membranes (47). A possible role for lipids in chemical communication is likewise documented (58). For instance, mating in reptiles may depend on epidermal skin lipids (59) and migratory behaviours in sea lamprey are regulated by larval-released fatty acid-derivatives (60). Bile, which contains phospholipids, mainly phosphatidylcholines (61), is metabolised in the liver and excreted into the environment to trigger chemical communication (62, 63). Lastly, in zebrafish embryos, these lipids may also originate from the yolk sac, being the most common membrane lipids (64). Since membrane lipids such as phosphatidylcholine are known not only as cellular signalling molecules, but also as conveying group identity between newly hatched catfish (*Plotosus lineatus*) (65), this supports a role for heat-mediated change in composition of membrane lipids as heat-stress related signalling molecules. For instance, *xkr8.2*, active within zebrafish lateral line hair cells (66) — which plays a role in chemical communication (67) — is overexpressed in response to stress metabolites. Its human orthologue *xkr8* flips phospholipids between the inner and outer membrane layers (68), so it could be used by receivers to interact with lipid stress metabolites. Further compounds functioning as stress metabolites could be peptides or proteins (69), which we could not characterise further. Additionally, several stress metabolites were matched on Metabolomics Workbench (70) with studies investigating stress, cancers, disorders, and infections, and/or in biofluids and excretions in several animal species, supporting their role as stress biomarkers and chemical signals.

As an outcome of this process, heat stress experienced by donors induced stress in naive receiver embryos. Molecular and phenotypic effects included accelerated development and impaired behavioural activity, as previously found for stress metabolites (16) and alarm cues (26). These phenotypic effects were similar to those induced by heat itself, but developmental acceleration caused by stress metabolites had a later onset at 4 dpf. Stress metabolites also initiated a suite of cellular signalling pathways different from those induced by heat. The most significantly downregulated gene in stress metabolite receivers was *si:ch211-214b16.2* (NOD2 ortholog) which is involved in intracellular signal transduction (71) and in the activation of apoptotic and immune pathways (72, 73). Further immune and apoptosis responses were induced via *fthl28* and *tlr18*. *tlr18* is a fish-specific toll-like receptor expressed in the skin, regulated by infection challenge and lipopolysaccharides (74), and may be homologous to the mammalian *tlr1* which binds lipopeptides (75). Therefore, *tlr18* may be directly responsive to lipid stress metabolites present in the medium, which we could independently confirm in group-exposed PET line zebrafish. Corroborating these results, brain transcriptomes of threespine stickleback *Gasterosteus aculeatus* exposed to predator and alarm odours were also enriched for apoptosis, immune, and signalling pathways (76). Moreover, several genes related to development such as keratins were activated by stress metabolites, which may explain the observed developmental acceleration in this treatment. In addition to hypoxanthine 3-N-oxide (C5H4N4O2 — for which we here found (C5H4N4O) to be more prevalent in the control condition), *Schreckstoff* is also known to contain other components such as extracellular polysaccharides (glycosaminoglycans), presumably but not exclusively chondroitin sulphate, and mucin proteins (77). The second major glycosaminoglycan in zebrafish is keratan sulphate (78), which is bound to proteoglycan, another component of the mucus layer next to mucins. In this contribution, we found that phosphocholine and sugars secreted from heat-stressed embryos significantly upregulated proteoglycan 4 (*prg4a*) and altered two transcripts involved in keratan sulphate biogenesis (*chs1* and *chst2b*) in receiver embryos, both in AB and PET line experiments. This lends support to proteoglycan 4 and keratan sulphate being signalling components alike to *Schreckstoff*, and since this was observed in receivers not senders of heat stress metabolites, it could mean that naive receivers may themselves propagate the release of *Schreckstoff-*like social anxiety cues to further embryos in a developing clutch. Of note, the complete knock-down of the chemosensory receptor *ora3* (olfactory receptor A like 3) in embryos group-exposed to SM suggests receptor depletion upon involvement in SM detection.

We can therefore conclude that heat effects have been propagated to receivers through stress metabolites, inducing similar phenotypic outcomes and molecular mechanisms related to growth acceleration, but also some characteristic differences to the direct stress response such as the absence of cortisol and the use of different signalling pathways (SI Appendix Fig. S13). Outside of the laboratory, high-density population scenarios such as developing fish clutches exposed to heat would experience stress metabolites simultaneously with heat, leading to combined exposure. Here, the combined treatment of heat and stress metabolites from heat-exposed embryos revealed that these larvae grew the most and also had the lowest behavioural activity. While heat was the predominant driver of this transcriptome, it also showed unique responses compared to the heat-only treatment, catalysed through the addition of stress metabolites. These involved differential expression of two chemosensory (vomeronasal receptor-like) genes, *ora3* and olfcb1, and genes involved in developmental acceleration via five genes in the apelin signalling pathway. Apelin signalling is a type of environmental signal processing with a wide array of physiological effects such as angiogenesis, renal fluid homeostasis, energy metabolism, immune response, embryonic development, and the neuroendocrine stress response (79), which hints at activation of the sympathetic nervous system SNS (80) and the emergence of chronic stress (81). We therefore anticipate that such positive stress feedback loops warrant further study due to the ever-increasing occurrences of diel and seasonal heat events.

## Acknowledgments

We acknowledge Alan Smith, Sonia Jennings, Emma Chapman, Robert Donnelly, and Paul Green for technical support. We are thankful to the animal facility staff of the University of Sheffield for providing us with a line of AB zebrafish. We thank all the members of the MolStressH2O and ChemEcolHull research clusters (University of Hull, UK) for valuable peer-review. We thank the members of the Metabolomics & Proteomics Facility of the University of York (including Tony Larson, Swen Langer, and Adam Dowle) for conducting the LC-MS/MS analysis and their valuable help in achieving the metabolomic project. The authors wished to acknowledge Matthew Arno and other members of the Edinburgh Genomics Facility for performing the RNA-sequencing and for their support in this project. We further acknowledge Chris Collins and the Viper High Performance Computing facility of the University of Hull and its support team.

## Ethics approval

All experiments were approved by the University of Hull Ethics committee under the approval U144b.

## Consent to participate

All authors consented to participation.

## Consent for publication

All authors consented to publication.

## Availability of data and material (data transparency)

All supplementary information, data, and analysis codes will be made publicly available upon manuscript acceptance.

